# Brain responses in aggression-prone individuals: A systematic review and meta-analysis of functional magnetic resonance imaging (fMRI) studies of anger- and aggression-eliciting tasks

**DOI:** 10.1101/2022.01.11.475895

**Authors:** Maja Nikolic, Patrizia Pezzoli, Natalia Jaworska, Michael C. Seto

## Abstract

**Background:** While reactive aggression (in response to a perceived threat or provocation) is part of humans’ adaptive behavioral repertoire, it can violate social and legal norms. Understanding brain function in individuals with high levels of reactive aggression as they process anger- and aggression-eliciting stimuli is critical for refining interventions. Three neurobiological models of reactive aggression–the limbic hyperactivity, prefrontal hypoactivity, and dysregulated limbic-prefrontal connectivity models–have been proposed. However, these models are based on neuroimaging studies involving mainly healthy individuals, leaving it unclear which model best describes brain function in aggression-prone individuals.

**Methods:** We conducted a systematic literature search (PubMed and Psycinfo) and Multilevel Kernel Density meta-analysis (MKDA) of nine functional magnetic resonance imaging (fMRI) studies of brain responses to tasks putatively eliciting anger and aggression in aggression-prone individuals alone, and relative to healthy controls.

**Results:** Aggression-prone individuals exhibited greater activity during reactive aggression relative to baseline in the superior temporal gyrus and in regions comprising the cognitive control and default mode networks (right posterior cingulate cortex, precentral gyrus, precuneus, right inferior frontal gyrus). Compared to healthy controls, aggression-prone individuals exhibited increased activity in limbic regions (left hippocampus, left amygdala, left parahippocampal gyrus) and temporal regions (superior, middle, inferior temporal gyrus), and reduced activity in occipital regions (left occipital cortex, left calcarine cortex).

**Conclusions:** These findings lend support to the limbic hyperactivity model and further indicate altered temporal and occipital activity in anger- and aggression-eliciting situations that involve face and speech processing.

## Introduction

Unplanned aggressive behavior that occurs following a perceived provocation is referred to as reactive aggression (1,2). It is distinct from proactive aggression, which is premeditated and instrumentally motivated (1,3). While reactive aggression is an adaptive human response in specific circumstances, it can also violate societal and legal norms. It accounts for more violent offenses than proactive aggression (4,5,6) and can have severe repercussions on victims (7).

High levels of reactive aggression can be a sign of emotional or cognitive problems that underlie socially inappropriate behavior, including a poor ability to regulate negative emotions (8) and poor executive functioning (9,10,11). High levels of reactive aggression can also be a symptom of personality and psychiatric disorders (12,13) as it is frequently observed in antisocial and Borderline Personality Disorders (BPD; 14,15), psychopathy (16) and Intermittent Explosive Disorder (IED; 17,18). Research on the factors that might predispose individuals to engage in reactive aggression is critical to advance interventions aimed at reducing this problem.

### Proposed Neurobiological Mechanisms of Reactive Aggression

Neuroimaging studies involving healthy non-aggressive individuals indicate that reactive aggression is associated with increased activation in the amygdala, which plays a pivotal role in processing emotionally-salient stimuli (19,20). In contrast, activity in the prefrontal cortex (PFC), involved in cognitive control, appears to be reduced (17,18,21). Additionally, reactive aggression has been linked with reduced limbic-PFC connectivity (22,23). Limbic-PFC connectivity is crucial for appropriate emotion regulation processes (24,25,26) as the orbitofrontal cortex (OFC) and dorsolateral prefrontal cortex (DLPFC) receive inputs from the amygdala and other medial temporal regions to integrate affective information (27,28). Reduced limbic-PFC connectivity may thus suggest deficits in regulating negative emotions (17,21,24,29).

Some neuroimaging studies examined differences in brain function between individuals with high levels of reactive aggression and non-aggressive controls. One study using functional magnetic resonance imaging (fMRI) examined emotional information processing and found that participants with IED exhibited greater amygdala activity, diminished OFC activity, and decreased connectivity between these regions during angry faces processing (17). Evidence of disrupted amygdala-OFC connectivity during the processing of angry faces in participants with IED (vs. healthy controls) has been replicated (18). In response to provocation, violent offenders (vs. non-aggressive controls) have been found to exhibit decreased connectivity between the left medial PFC and left amygdala (29), greater activity in the amygdala and striatum, and reduced amygdala-PFC and striatal-PFC connectivity (21). Altered activity in these networks during reactive aggression might reflect poor attenuation of emotional responses (22,23,24).

Based on a qualitative review of prior neuroimaging studies, reactive aggression appears to be associated with amygdala hyperactivity (17,18,21), PFC hypoactivity (17, 30, 31), and dysregulated limbic-PFC connectivity (17,21,29). However, a systematic review supported altered limbic-PFC connectivity but not amygdala hyperactivity, and PFC hypoactivity (32). Because this review only included studies involving healthy participants, however, it remains unclear what activation patterns would exist in aggression-prone individuals. Another review (33) reported two meta-analyses: one including studies of cognitive tasks in aggressive psychiatric individuals, and one including studies of aggression-eliciting tasks in healthy non-aggressive individuals. The first meta-analysis found reduced activity in the precuneus, a region involved in cognitive function (34), in aggressive psychiatric individuals (vs. controls). The second found activation in the right postcentral gyrus during aggression-eliciting tasks in healthy non-aggressive individuals, but no activation in regions associated with emotion generation and regulation (e.g., amygdala and PFC). This second analysis examined brain activity in individuals with high trait aggression during executive function tasks, but not during potentially aggression-eliciting or emotionally salient paradigms that can elicit reactive aggression. Moreover, this analysis did not differentiate reactive and proactive aggression.The present systematic review addresses these gaps by investigating brain activity patterns in individuals with a history of antisocial behavior relative to healthy controls during potentially aggression-eliciting paradigms.

## Methods and Materials

To promote transparency and minimize risk of bias, we pre-registered our protocol on PROSPERO on February 2, 2021 (CRD42021211242).

### Study Selection

We conducted our systematic review (January 30^th^-March 18^th^, 2021) using PubMed (Medline) and APA Psycinfo (Ovid) advanced search builder. The following search criteria and keywords were used: (*“aggression”* or *“aggressive”*) with (*“reactive”* or *“expressive”* or *“hostile”* or *“impulsive”* or *“violent”* or *“explosive” or “anger”* or *“angry”* or *“overt”* or *“emotional”*) paired with (*“fMRI[tw]”* or *“functional magnetic resonance imaging”*), for entries dating January, 1990-March 18^th^, 2021. We used Covidence (https://www.covidence.org) to organize, manage, and detect duplicate citations. We followed the Preferred Reporting Items for Systematic Reviews and Meta-Analyses (PRISMA) guidelines for systematic review (35). Two authors (MN, PP) reviewed and screened titles, abstracts, and full texts according to pre-defined selection criteria, and independently coded information on included sources in a data extraction matrix. Conflicts were resolved through discussion and involving other authors (MCS, NJ). The number of records identified, included, and excluded in the process are depicted in **Figure 1**.

**Figure 1.**
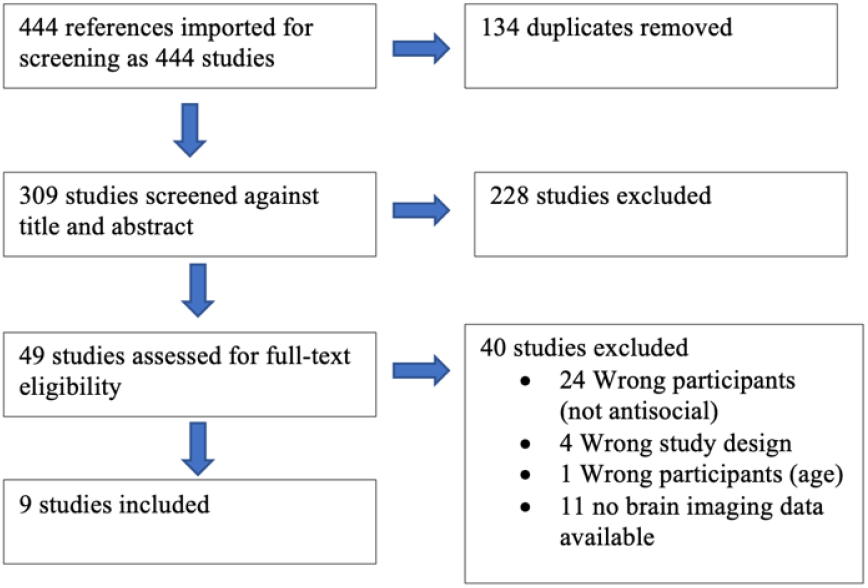
PRISMA flowchart illustrating the literature search and study selection process.

### Inclusion/Exclusion Criteria

We included studies that met the following criteria:

1. Peer reviewed, in English;
2. Reported original data from participants aged ≧16yr;
3. Reported whole-brain thresholded results in a standard anatomical space. Studies that used only a region-of-interest approach were excluded;
4. Examined brain activity during aggression-eliciting paradigms, namely, paradigms that have been found to evoke reactive aggression outside the scanner and/or to be a valid proxy for reactive aggression in the scanner. Studies using resting-state paradigms and/or cognitive paradigms with no anger or aggression-eliciting component were excluded;
5. Involved participants with high levels of reactive aggression and a non-aggressive control group. Studies involving only non-aggressive participants were excluded.

We considered participants as aggression-prone if they met at least one of the following criteria:

1. Have a Diagnostic and Statistical Manual of Mental Disorders (36) or International Classification of Diseases (37) psychiatric diagnosis that specifies patterns of behavior disregarding and violating the rights of others (e.g., antisocial personality disorder, IED, conduct disorder) with documented history of overtly harming others; and/or
2. Have been charged with, convicted, or incarcerated for aggressive behavior, against persons or property; and/or
3. Scored above a normative threshold on a standardized behavioral assessment of aggression, whether clinician-rated or standardized psychometric measures (e.g., Buss and Perry Aggression Questionnaire; 38).

### Data Synthesis

A quantitative analysis was conducted to compare brain activation during aggression-eliciting paradigms in aggression-prone individuals and controls. This included stratifying brain activation patterns according to condition (aggression-eliciting vs. control) and participant group (aggressive vs. controls), and conducting a coordinate based meta-analysis.

#### Coordinate Based Meta-Analysis: Multi-level Kernel Density Analysis

We conducted a multilevel kernel density analysis (MKDA), which summarizes evidence of activation from the included studies for X (left-right), Y(posterior-anterior), and Z (inferior-superior) peak coordinates in Montreal Neurological Institute (MNI) space (39). We chose this method because the alternative, Activation Likelihood Estimation (ALE), treats all peak coordinates across studies as independent units of analysis and, thus, increases the risk of the results being driven by a subset of studies showing the same activation peaks (40).

In MKDA, the unit of analysis is the sample size-weighted proportion of studies that report activation differences in a spatial location, which increases generalizability. MKDA summarizes evidence for activation in a local neighborhood around each voxel in a standard brain atlas, and reports coordinates in reference to a statistical contrast map (SCM) of activated brain regions for each study (39). Consistency and specificity across studies is analyzed in the neighborhood of each voxel, and consistency is determined by how many SCMs are activated near a voxel (41). A 3D histogram of peak locations is constructed and smoothed with a spherical indicator function of radius; this convolution occurs within each SCM and results in the creation of contrast indicator maps (CIMs). Weighted CIMs are thresholded based on the maximum proportion of activated comparison maps under the null hypothesis distribution, where the distribution of contiguous regions of activation in the CIMs are randomly and uniformly distributed throughout the brain (**Figure 2**). Sample size is also considered, as the precision of the estimates from each study is proportional to the square root of the study sample size (39). Lastly, 5000 Monte Carlo simulations were performed to compare the observed density map to a null distribution of density maps created by identifying clusters of activated voxels for each SCM and then randomizing cluster centers within gray matter in the standard brain.

**Figure 2.**
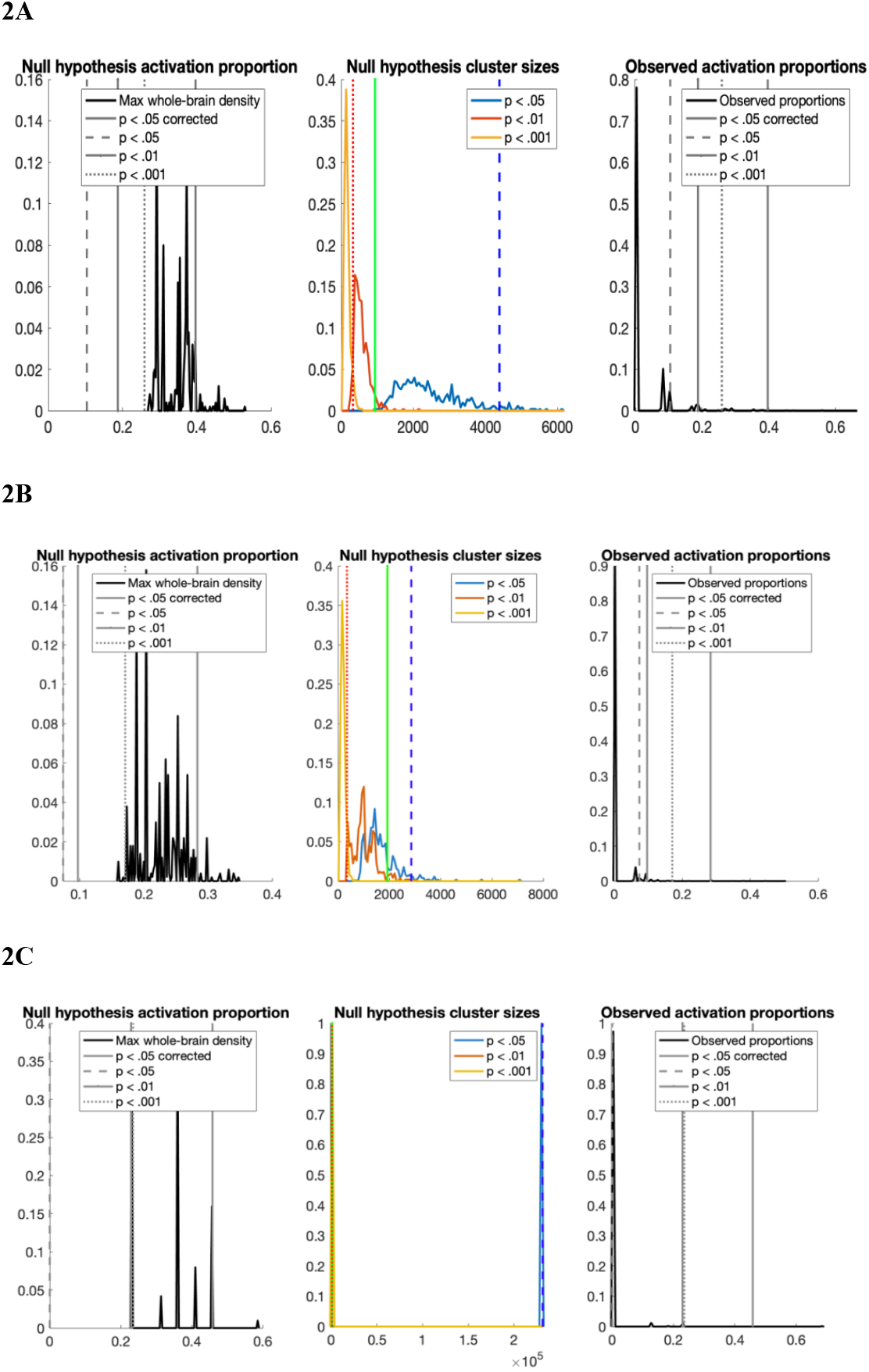
Monte Carlo null hypothesis activation proportion and thresholding for aggression-prone individuals (2A), aggression-prone individuals vs. healthy controls (2B) and healthy controls vs. aggression-prone individuals (2C). Null hypothesis activation proportion: The weighted proportions of comparison, namely the maximum proportion of activated comparison maps under the null hypothesis, are plotted on the x axis. Null hypothesis cluster sizes: The largest cluster of contiguous voxels, which is important for extent-based thresholding, is reported on the x axis. Observed activation proportions: the maximum density value (height threshold) across all studies following each Monte Carlo iteration is presented on the x axis. In Figure 2A, p<0.05 MKDA-height corrected represents a height threshold of 0.40 and includes 251 voxels. p<0.001 represents a height threshold of 0.26 and extent threshold of 324 (includes 3592 significant voxels). p<0.01 represents a height threshold of 0.19 and extent threshold of 936 (includes 2350 significant voxels). p<0.05 represents a height threshold of 0.10 and extent threshold of 4386 (includes 0 significant voxels). In Figure 2B, p<.05 MKDA height-corrected represents a height threshold of .28 and includes 1030 voxels; p<0.001 represents a height threshold of .17 and extent threshold of 350 (includes 1104 significant voxels); p<.01 represents a height threshold of .10 and extent threshold of 1926 (includes 0 significant voxels); p<.05 represents a height threshold of .07 and extent threshold of 2858 (includes 0 significant voxels). In Figure 2C, p<.05 MKDA height-corrected represents a height threshold of .46 and includes 369 voxels; p<.001 represents a height threshold of .24 and extent threshold of 896 (0 significant voxels); p<0.01 represents a height threshold of .23 and extent threshold of 902 (0 significant voxels); p<0.05 represents a height threshold of 0 and extent threshold of 231202 (0 significant voxels).

##### Software, Data, and Code Availability

We performed analyses in MATLAB v.R2021a using the MKDA toolbox developed by Tor Wager (https://github.com/canlab/CanlabCore). Our code is publicly available on the Open Science Framework (http://osf.io/CG94W). To identify activated brain regions associated with the generated MNI coordinates, we used Neurosynth (https://neurosynth.org/locations/).

## Results

### Participants

Our systematic literature review included 9 studies involving 230 aggression-prone individuals and 235 healthy controls. Tables 1–3 report information on demographic and clinical characteristics (**Tables 1** & **2**), offense histories, and antisocial behavior assessments (**Table 3**).

**Table 1.**
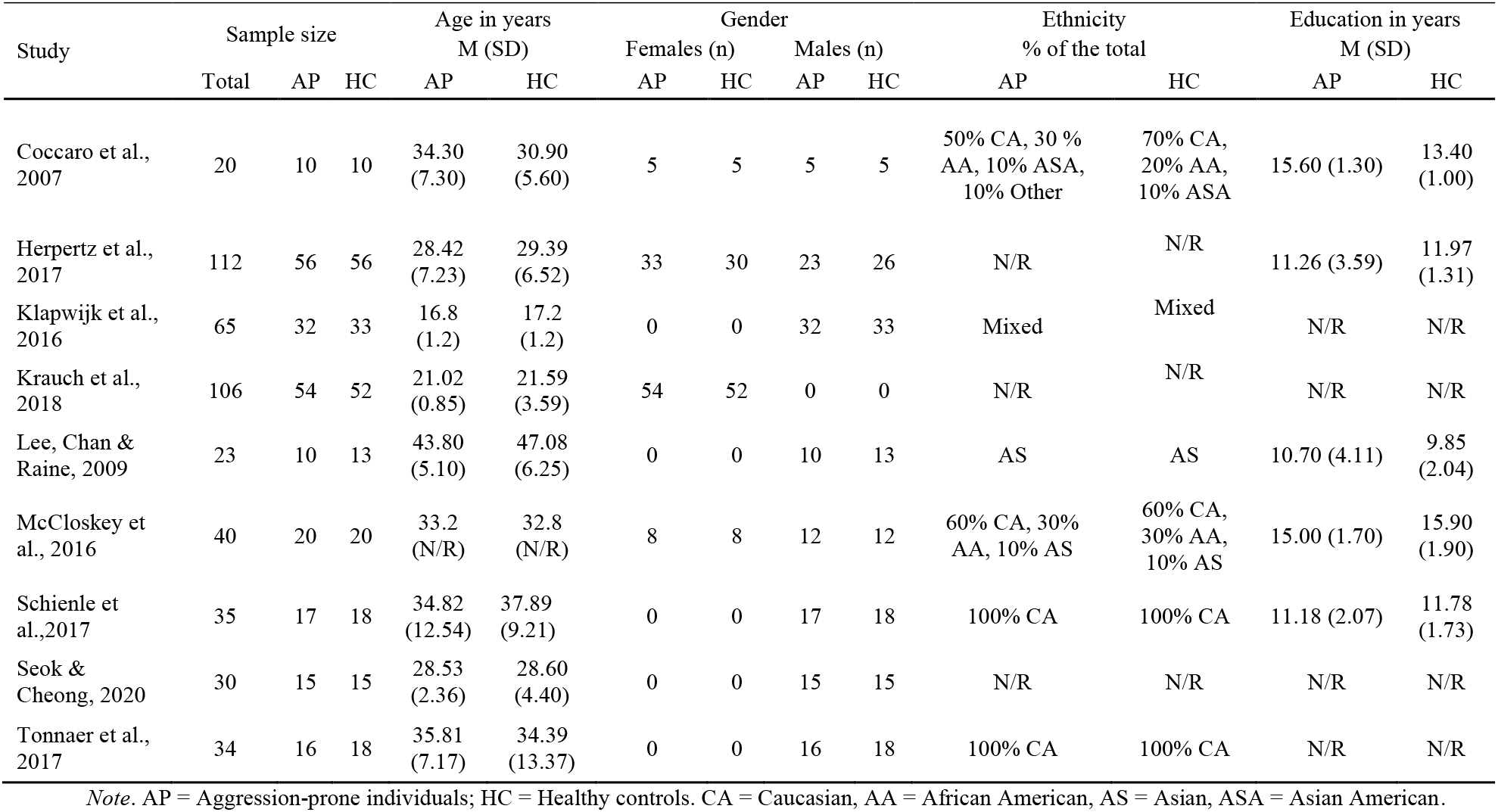
Demographic characteristics of participants

**Table 2.**
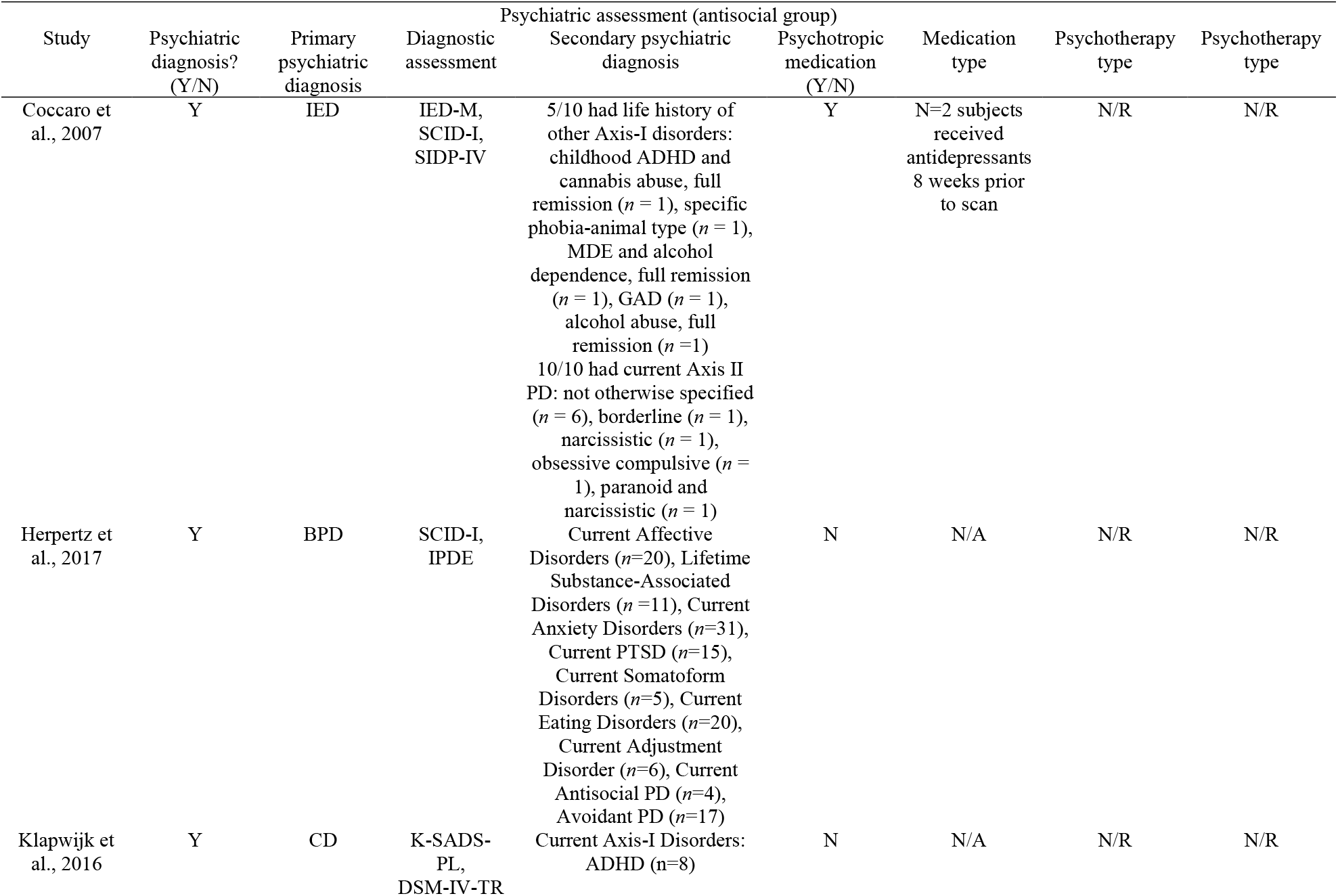
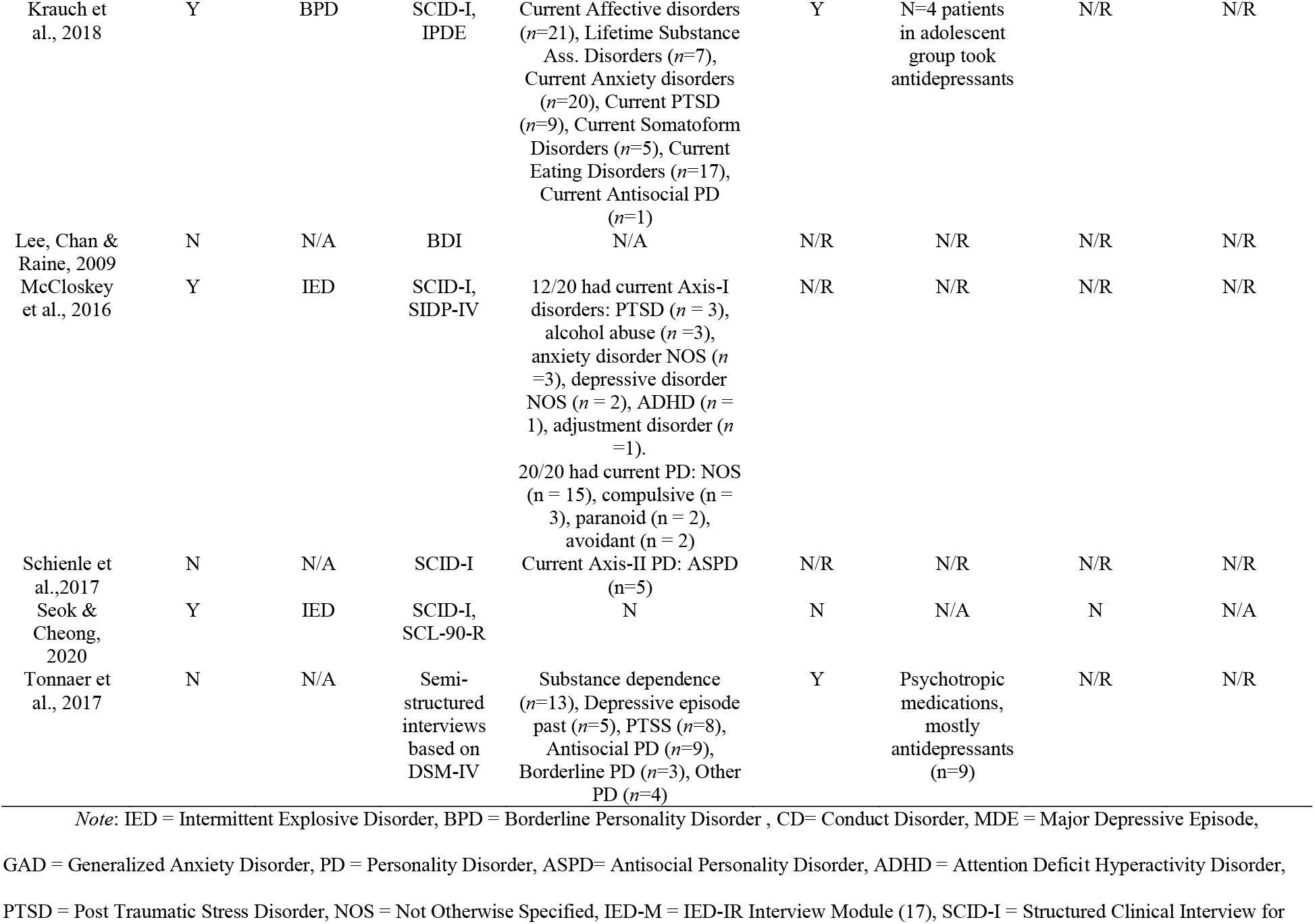
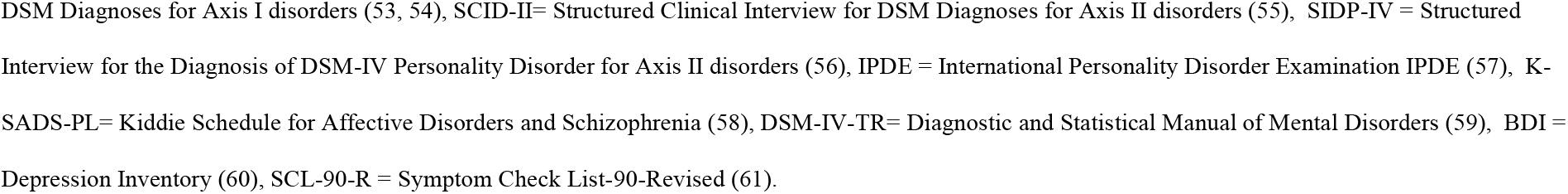
Clinical characteristics of participants

**Table 3.**
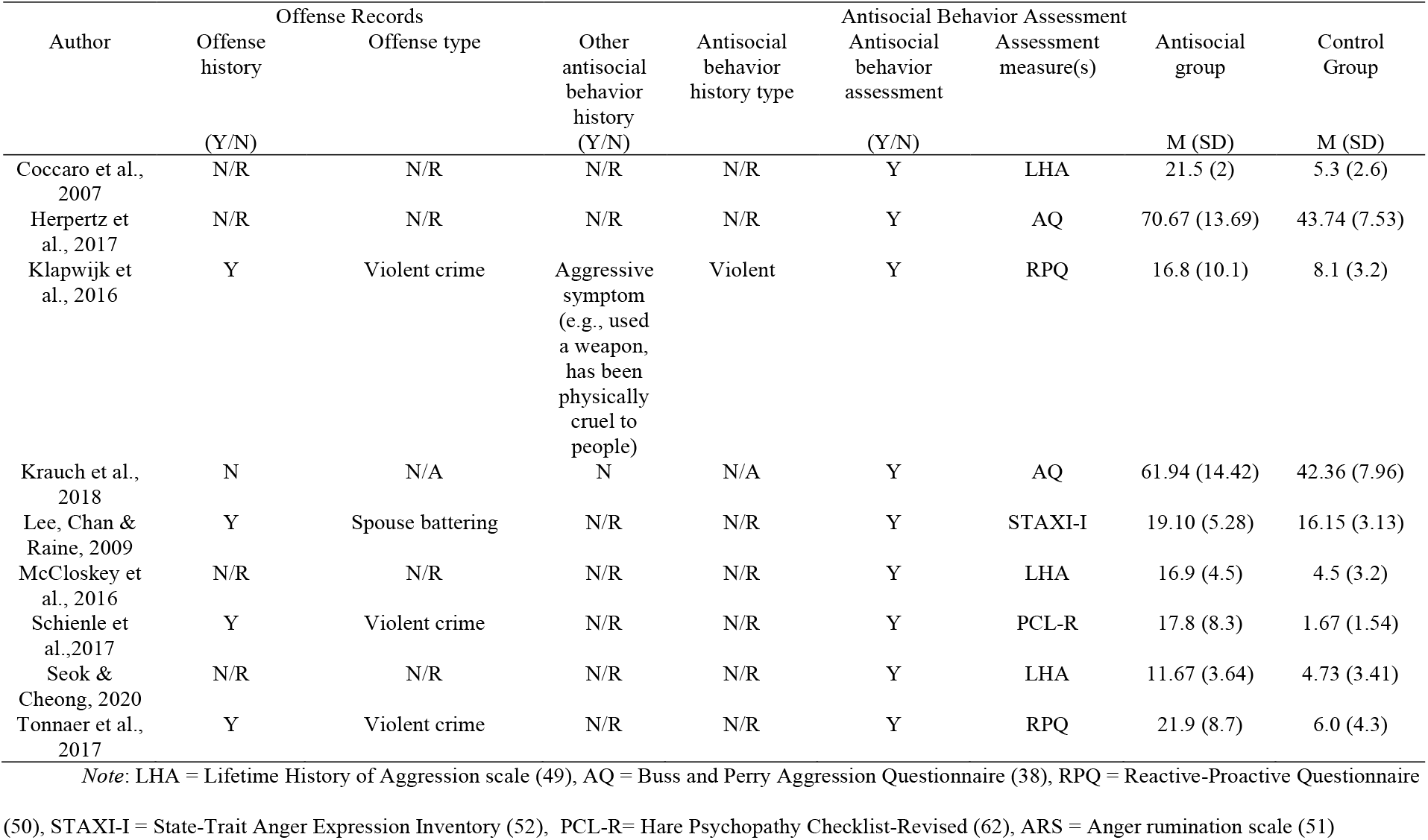
Offense histories and assessments of antisocial behavior

Aggression-prone and control groups were matched for demographic characteristics. Aggression-prone individuals were aged 17-44yr, healthy controls were 17-47yr. All participants completed secondary education, except for one study which included adolescents (42). Three studies involved both females and males (17,18,43), five involved males only (42,44,45,46,47), and one was comprised of females only (48). Half reported ethnicity, with ∼70% of participants being Caucasian (17,18,45,47), while one included only Asian participants (44).

Seven out of nine studies involved aggression-prone individuals with a psychiatric disorder, including: BPD (43,48), IED (17,18,46), Conduct Disorder (42), or multiple disorders (47). Seven studies reported information on comorbid conditions, which included personality disorders, affective disorders, and substance disorders (17,18,42,43,45,47,48). Three studies reported that some participants were receiving psychotropic medication (antidepressants, most commonly; 17, 47, 48).

Four studies reported that participants had criminal offense histories, including attempted manslaughter or murder, assault, or domestic violence (42, 44, 45, 47). An assessment of antisocial behavior was conducted in each of the included studies, by means of the Lifetime History of Aggression (49), Buss and Perry Aggression Questionnaire (38), Reactive-Proactive Questionnaire (50), Anger Rumination Scale (51), or State-Trait Anger Expression Inventory (52). Aggression-prone participants scored higher than normative values on all these measures, and significantly higher than their respective control groups.

### Tasks

fMRI tasks aimed at eliciting aggression included script-driven-imagery tasks (43,48,47), a personal-space intrusion task (45), an anger-eliciting task (42), and viewing emotional images (17,18,44) and videos (46). During script-driven-imagery tasks, participants listened to audiotapes and were asked to imagine the scenes as vividly as possible to provoke an intense emotional response. We compared conditions that likely elicited aggression (e.g., “anger induction”, “anger engagement”) to neutral conditions. In the personal-space intrusion task, participants viewed static or “approaching” neutral faces, where pictures were enlarged to the point where only the mouth and eyes were visible, creating the impression of an invasion of personal space (45). We compared the approaching condition to the static face condition. In the anger-eliciting task, participants read the responses of a fictional opponent following an unfair distribution of tokens (42). We compared angry responses (potentially aggression-eliciting) to happy responses (baseline). In the implicit emotion processing task, participants viewed faces expressing emotions and had to identify their gender (17). We compared the angry face viewing condition (potentially aggression-eliciting) to rest (blank screen; no neutral face condition available). In the explicit emotion processing task, participants had to identify the emotional valence of neutral, positive, or negative faces; we compared angry vs. neutral faces. During the passive viewing of images tasks, participants viewed neutral, positive, and aggressive pictures, the latter involving violent pictures with female victims or general aggressive threats (e.g., person pointing a gun). The task involving passive viewing of video clips included anger-related and neutral clips; for these, we contrasted the anger-/aggression-related and neutral conditions.

### Multi-Level Kernel Density Analysis (MKDA) Results

We considered significant voxels to be those >95^th^ percentile value under the null hypothesis (threshold derived from Monte Carlo simulations) (63). This generated the final regions of activated contrast indicator maps. Herpertz et al. (2017) had the largest sample size (N=112) and thus the greatest influence on the analysis.

**Figure 3** depicts the final map of significant results generated after the weighted average of the contrast indicator maps was compared and thresholded with the maximum proportion of activated comparison maps under the null hypothesis distribution. Peak brain activations generated for within-group (**Table 5**) and between-group (**Tables 6** & **7)** analyses are denoted.

**Table 4.**
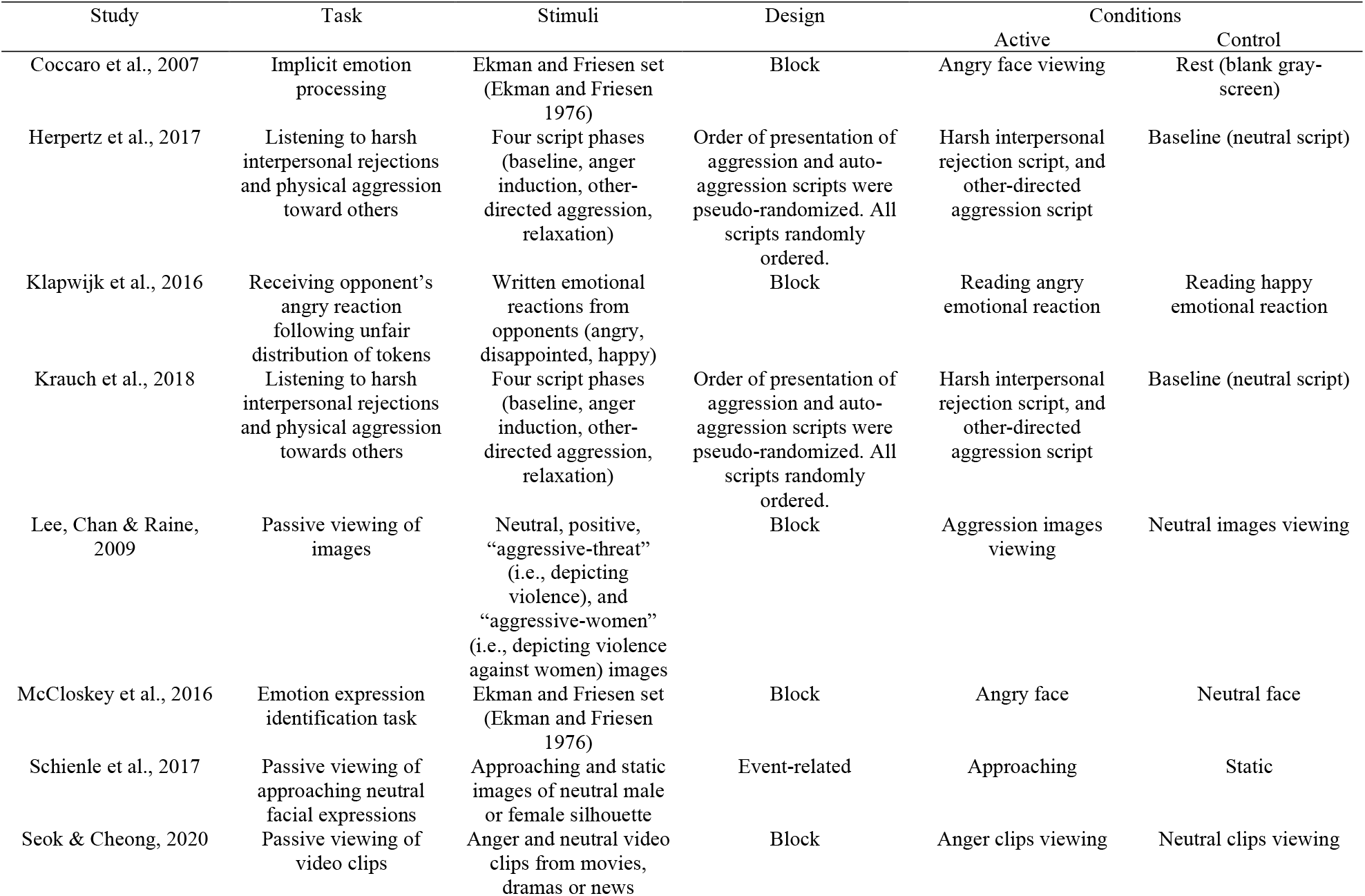
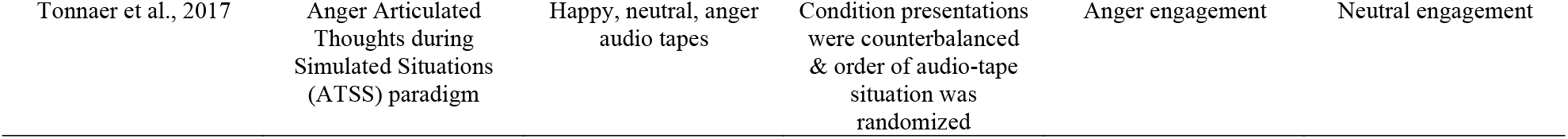
Functional Magnetic Resonance Imaging (fMRI) tasks

**Table 5.** Peak activations in aggression-prone participants during aggression-eliciting relative to control conditions

| Cluster | x | y | z | k (voxels) | Brain region |
| --- | --- | --- | --- | --- | --- |
| A) Overall analysis |  |  |  |  |  |
| 1 | 56 | -18 | -1 | 4 | Right superior temporal gyrus (64) |
| 2 | 58 | -6 | 0 | 2 | Right superior temporal gyrus/superior temporal sulcus (65) |
| 3 | 14 | -92 | 4 | 1 | Occipital pole (66) |
| 4 | 14 | -88 | 12 | 26 | Posterior cingulate cortex (67) |
| 5 | 50 | -2 | 46 | 218 | Right precentral gyrus (68) |
| B) Additional regions at extent threshold: stringent and size $\geq 10$ | | | | | |
| 1 | -60 | -18 | 4 | 378 | Left superior temporal gyrus (69) |
| 2 | 48 | 16 | 24 | 425 | Right inferior frontal gyrus (70) |
| 3 | 10 | -58 | 44 | 411 | Right precuneus (71) |
| 4 | 8 | 14 | 66 | 342 | Right supplementary motor area (SMA) (72) |
| C) Additional regions at extent threshold: medium and size $\geq 10$ | | | | | |
| 1 | 16 | -68 | 28 | 180 | Right posterior cingulate cortex (73) |
| 2 | 44 | 14 | 36 | 81 | Right middle frontal gyrus (74) |
| 3 | -2 | -54 | 54 | 445 | Precuneus (75) |

**Table 6.** Peak activations in aggression-prone participants>healthy controls during aggression-eliciting relative to control conditions

| Cluster | x | y | z | k (voxels) | Brain region |
| --- | --- | --- | --- | --- | --- |
| A) Overall analysis |  |  |  |  |  |
| 1 | -30 | -12 | -26 | 515 | Left hippocampus/amygdala (76) |
| “ | -32 | -16 | -30 | 121 | Left parahippocampal gyrus (77) |
| “ | -28 | -8 | -28 | 178 | Left hippocampus/amygdala (78) |
| “ | -30 | -14 | -22 | 216 | Left parahippocampal gyrus (79) |
| 2 | 52 | 2 | -22 | 515 | Right superior temporal sulcus/middle temporal gyrus (80) |
| “ | 52 | 2 | -28 | 100 | Right middle temporal gyrus/superior temporal sulcus (81) |
| “ | 52 | -4 | -20 | 104 | Right middle temporal gyrus (82) |
| “ | 58 | 2 | -22 | 79 | Right middle temporal gyrus and inferior temporal gyrus (83) |
| “ | 50 | 4 | -20 | 232 | Right middle temporal gyrus (84) |
| B) Additional regions at extent threshold: stringent and size $\geq 10$ | | | | | |
| 1 | -26 | -2 | -24 | 74 | Left amygdala (85) |

**Figure 3.**
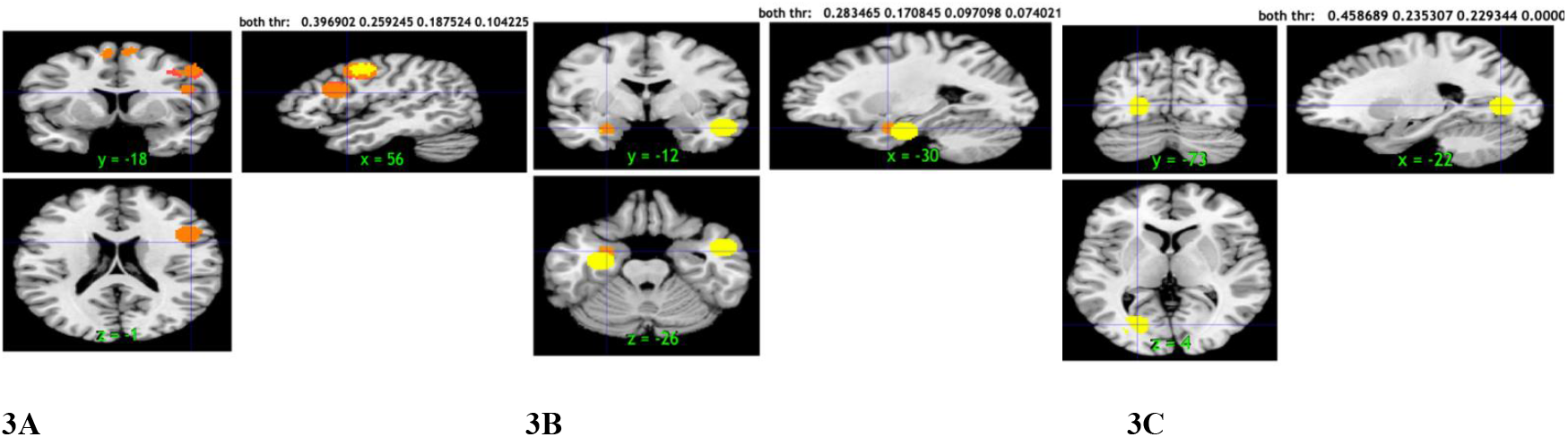
Proportion of activated CIMs (significant regions) in aggression-eliciting protocols>control conditions in aggression-prone individuals (3A), aggression-prone individuals relative to healthy controls (3B) and in healthy controls relative to aggression-prone individuals (3C). Regions depicted in yellow: significant at p<.05 MKDA-height corrected (representing threshold 0.40 with 251 voxels in 3A; threshold 0.28 with 1030 voxels in 3B and threshold 0.46 with 369 voxels in 3C). In those regions, the proportion of SCMs activated within r mm of the voxel was greater than would be expected by chance. Regions depicted in orange: significant at p<.05 cluster-extent corrected with primary alpha levels of .001 (representing threshold 0.26 with 3592 significant voxels in 3A; threshold .17 with 1104 significant voxels in 3B. 3C has no orange regions because there are 0 significant voxels at threshold 0.24. Those regions were large enough in size to expect that we would only see such a cluster in the brain by chance 5% of the time (41, 63). In those regions, MKDA was extent-based thresholded, meaning that the largest set of contiguous voxels was saved at each Monte Carlo simulation, and cluster extent threshold value was determined as the 95^th^ percentile of these values across each iteration (in this case: 1, 324, 936, 4386 for 3A; 1, 350, 1926, 2858 for 3B, and 1, 896, 902, 231202 for 3C; 63).

Within-group MKDA indicated multiple regions that activated more in the aggression-inducing vs. control conditions in aggression-prone individuals, comprising a total of 2513 voxels (**Table 5**). These regions included the right/left superior temporal gyrus, right inferior frontal gyrus, right middle frontal gyrus, right precentral gyrus, occipital pole, SMA, posterior cingulate cortex (PCC) and precuneus. The clusters are shown in **Figure 3A**, with coordinates (56, −18 −1) representing the first cluster.

Between-group MKDA indicated sets of regions that were more strongly activated in aggression-prone vs. control participants, comprising a total of 2134 voxels (**Table 6**). These included the left hippocampus, left amygdala, left parahippocampal gyrus, right superior temporal sulcus, right inferior temporal gyrus and middle temporal gyrus. The clusters are shown in **Figure 3B**, with coordinates (−30, −12, −26) representing the first set of clusters. Between-group MKDA also indicated regions that were more strongly activated in healthy controls vs. aggression-prone participants, comprising a total of 727 voxels (**Table 7**). These regions included the left occipital cortex, left medial occipital cortex, left lingual gyrus of the occipital lobe, left calcarine cortex and left V2. The clusters are shown in **Figure 3C**, with coordinates (−22, −73, 4) representing the first set of clusters.

**Table 7.**
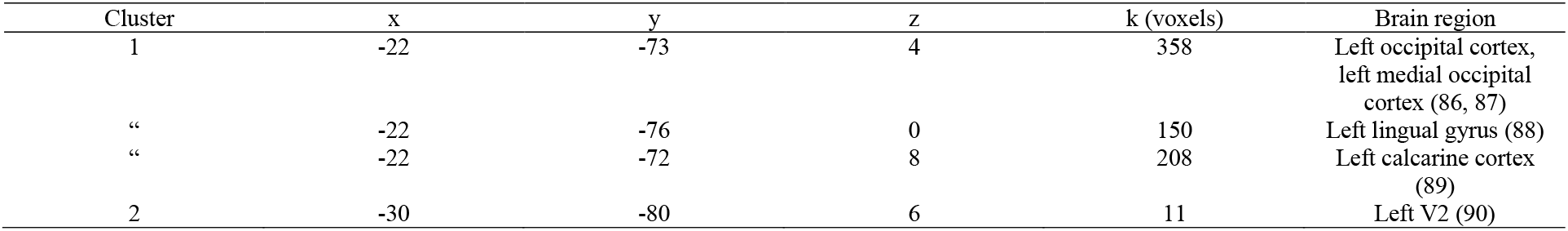
Peak activations in healthy controls>aggression-prone participants during aggression-eliciting relative to control conditions (overall analysis)

| Cluster | x | y | z | k (voxels) | Brain region |
| --- | --- | --- | --- | --- | --- |
| 1 | -22 | -73 | 4 | 358 | Left occipital cortex,<br>left medial occipital<br>cortex (86, 87) |
| “ | -22 | -76 | 0 | 150 | Left lingual gyrus (88) |
| “ | -22 | -72 | 8 | 208 | Left calcarine cortex<br>(89) |
| 2 | -30 | -80 | 6 | 11 | Left V2 (90) |

## Discussion

### Neural Responses to Elicited Aggression in Aggression-Prone Individuals

Our within-group MKDA analysis included five studies (18,42,43,45,48) and indicated that aggression-prone individuals exhibited greater activation in several brain regions, including the superior temporal gyrus, right inferior frontal gyrus, right PCC and precuneus, during aggression-eliciting compared to control conditions.

The temporal lobe is involved in speech, sound, and visual affect processing (91,92,93). In particular, the superior temporal gyrus is involved in social cognition (94) through its role in processing auditory information (e.g., spoken words; 95) and visual information (e.g., eye and body movements; 93, 96). Consistent with this, our results indicated that the superior temporal gyrus was activated when aggression-prone individuals processed visual and auditory aggression-eliciting stimuli. The superior temporal gyrus is one of the principal sites of high-level sensory information convergence (97). Since social information processing might be compromised in aggression-prone individuals (98), who frequently display a hostile attribution bias (99,100), superior temporal gyrus activation might reflect a neural hypersensitivity to anger-provoking social cues (44,101,102). Given evidence of direct projections from the superior temporal gyrus to the hippocampal entorhinal cortex (103), this may reflect the acquisition of new emotionally-salient memories (104,105).

The right inferior frontal gyrus (IFG) is part of the PFC and is an essential component of response inhibition (106,107,108, 109, 110). Since aggression has been linked to response inhibition deficits (111,112), enhanced right IFG activity in aggression-prone individuals might reflect an effort to inhibit aggressive responses, which is particularly relevant as participants had to presumably refrain from exhibiting aggressive behaviors in the scanner. Studies involving healthy individuals have shown that enhancing right IFG activity through brain stimulation improves inhibition in a stop signal task (113). Future research could explore whether stimulating this area enhances response inhibition in aggression-prone individuals.

The PCC, middle temporal gyrus and precuneus are part of the default mode network (DMN; 114,115) and represent some of the most metabolically active brain regions at-rest and during cognitive tasks (116,117). The PCC integrates information across cortical networks (118), supporting behavioral regulation in changing environments (119,120). The precuneus is a regulatory region strongly interconnected with the fronto-parietal network (FPN;34), and its disrupted activation has been observed in high trait aggressive individuals (33). Increased activation of the PCC and precuneus in aggression-prone individuals may thus reflect ineffective DMN suppression, or an increased effort to suppress behavioral responses to provocation.

### Differences between Aggression-Prone Individuals and Healthy Controls

Our between-group MKDA analysis included 8 studies (17,18,42,43,44,46,47,48) and revealed greater limbic activation (left amygdala, left hippocampus, left parahippocampal gyrus), greater temporal activation (superior, middle and inferior temporal gyrus), and decreased occipital activation (left occipital cortex and left calcarine cortex) during aggression-eliciting conditions (vs. baseline) in aggression-prone individuals relative to healthy controls.

#### Expanding the Amygdala Hyperactivity Model of Reactive Aggression

The finding of increased limbic activation provides support and extends the amygdala hyperactivity model of reactive aggression (22,23). This is in contrast with a previous meta-analysis (33), which did not find abnormalities in limbic activation in individuals with high trait aggression. However, this discrepancy might be due to the inclusion of cognitive tasks (2); here, we focused on emotion generation and regulation to provocation (or, perceived provocation).

Our results further expand an amygdala hyperactivity model of reactive aggression by suggesting that other limbic areas that have received less attention, namely the hippocampus and parahippocampal gyrus, are also implicated. The hippocampus is embedded in the temporal lobe, on the posterior part of the limbic lobe (121) and is mainly involved in episodic memory (122,123) but also in emotion regulation and emotional memory processing (124,125). Structural abnormalities in the hippocampus, such as exaggerated asymmetry, have been associated with impulsive and disinhibited behaviors (126). Disruption of the PFC-hippocampal circuitry has been associated with affect dysregulation and impulsive disinhibited behavior (126,127,128). Therefore, the hippocampus may play a role in modulating aggressive responses (128). Our findings support this possibility and motivate further work on the role of the hippocampus in reactive aggression. For instance, enhanced hippocampus and parahippocampal gyrus activation in aggression-prone individuals during provocation might reflect enhanced recall of associative memories related to aggression (129).

Our results further point to the importance of amygdala-hippocampal coupling (130). These regions display bi-directional functional relationships during encoding of emotional events (130) and contribute to forming semantic representations of emotionally-valanced stimuli (131,132). The hippocampus forms episodic representations of the emotional significance of events, which influence amygdala responses when emotional stimuli are next encountered (133).

Our findings also suggested left hemispheric lateralization in aggression. It has been postulated that the left amygdala is involved in specific and sustained stimulus evaluation, whereas the right amygdala is engaged in the automatic detection of emotional stimuli (134,135). Thus, it is possible that aggression-prone individual engaged in a more detailed analysis of the aggression-eliciting stimuli. Abnormalities in left amygdala-hippocampal coupling have been associated with deficits in the perception of social cues, including facial expressions (136). In particular, a meta-analysis found increased left amygdala and left hippocampus activation during the processing of negative emotional stimuli in those with BPD (136). Since most aggression-prone individuals in our meta-analysis had psychiatric disorders, the enhanced amygdala-hippocampus co-activation might also be a distinctive feature of reactive aggression in the context of psychiatric conditions, but this speculation warrants further study.

#### No Support for the Limbic-Prefrontal Models of Reactive Aggression

Our results did not provide direct support for the PFC hypoactivity model of reactive aggression (17,18,22). However, we did not perform functional connectivity analyses, and thus did not test models that emphasize effective interactions between amygdala and PFC regions in the control of aggressive behavior (17). Limbic-PFC models conceptualize reactive aggression as a failure in top-down control systems (i.e., PFC) to inhibit aggressive reactions generated in the limbic system (23). Consistent with this, prior studies observed amygdala hyperactivity and PFC hypoactivity in violent offenders (21,137), and decreased attenuation of amygdala reactivity by the PFC in aggression-prone individuals with BPD (136,138). Since our aggression-prone participants displayed abnormal occipital activation (see more below), one possibility is that PFC areas influenced amygdala activity indirectly, through modulation of other regions directly connected to it, namely perceptual areas in the occipital cortex (139). Indeed, regions like the DLPFC, implicated in the reappraisal of negative emotion through attenuation of amygdala responses (140,141), have sparse direct connections to the amygdala (142). Future functional connectivity studies are needed to elucidate this possibility.

#### Novel Findings: Role of the Temporal and Occipital Regions in Reactive Aggression

Our finding of greater activation in the superior temporal gyrus, right inferior temporal gyrus and middle temporal gyrus in aggression-prone individuals (143) is novel. The temporal lobe is involved in speech, sound and facial processing (91, 92, 93). The inferior and middle temporal gyri are mainly associated with visual perception and multimodal sensory integration (144,145) but also emotional face processing (146). Damage to these regions has been related to deficits in tasks requiring visual discrimination and recognition (147). The activation of these areas seems intuitive given that half of our studies involved viewing faces or videos, and half involved listening to anger-eliciting scripts or audiotapes.

To our knowledge, decreased activation in the left occipital cortex in aggression-prone individuals has not been previously noted. The occipital lobe is the visual processing area of the brain; it is implicated in object and face recognition, visuospatial processing, and visual memory formation (148). Prior research indicated that the amygdala facilitates perception and attention to emotionally salient stimuli through projections to the visual stream (149), and by modulating activation in higher sensory processing areas (150). Our results suggest that the mechanism by which the amygdala facilitates perception of aggression-eliciting stimuli through its modulation of visual streams might be dysregulated in aggression-prone individuals.

### Limitations

Our meta-analysis is limited by the possibility of publication bias (and, thus, false positive results;151,152) and by the small number of studies, which precluded examining potential moderators such as task type. Studies were characterized by small samples (potentially under-powered) and substantial variation in analyses parameters (153). MKDA adjusts for some of these biases by ensuring that larger and more rigorous studies exert the highest effects. Meta-analytic methods including small-sample adjustments, profile likelihood, or hierarchical Bayesian models (154) are recommended. Furthermore, included participants were 70% white, which challenges generalizability to other groups.

The included studies were heterogenous in how they defined aggression proneness. Four studies included violent offenders, some of whom were diagnosed with Antisocial Personality Disorder (45, 47), Conduct Disorder (42), or no diagnoses (44). Although these participants committed a violent crime, we cannot be certain that they all engaged in reactive aggression following a provocation, as their violent behavior may have been premeditated. For example, antisocial personality traits predict both higher reactive and proactive aggression (155). It would be beneficial if studies reported systematic accounts of participants’ reactive aggression history.

Paradigms used in fMRI research to study reactive aggression are heterogeneous, and target proxies of reactive aggression, rather than reactive aggression *per se;* as such, they likely lack external validity and experimental realism (156). While reactive aggression is incompatible with the scanning environment and, therefore, difficult to measure, novel protocols should be developed to effectively study the neural basis of reactive aggression. For example, more realistic tasks could combine virtual reality (VR) and fMRI (157,158). Exploring the use of VR to elicit aggression will require specific safety protocols to minimize ethical and safety concerns (159). Alternative devices that are less sensitive to motion, such as electroencephalogram (EEG) or functional near-infrared spectroscopy (fNIRS) could also be used (160,161).

## Conclusions

These findings lend support to the limbic hyperactivity model and further implicate altered temporal and occipital activity in anger- and aggression-eliciting situations that involve face and speech processing. Future studies can advance our understanding of reactive aggression by examining potential participant- and task-related moderators, functional connectivity, and specific hypotheses derived from our findings. For example, given our implication of the hippocampus, future studies could examine differences between aggression-prone and healthy individuals in mnemonic performance while processing aggression-eliciting stimuli.

## Acknowledgments

Scripts available at osf.io/CG94W. Authors have no conflicts of interest to disclose.

## Notes

**Sources of financial support:** Oversight Fund of the Integrated Forensic Program at the Royal Ottawa Health Care Group, Ottawa, Ontario, Canada (PP). The funding sources had no role in the preparation of this manuscript or the decision to submit it for publication.

https://osf.io/cg94w/

### Competing Interest Statement

The authors have declared no competing interest.

